# Neuronal Grin2c expression and the cocaine engram: a medial prefrontal cortex substrate for conditioned place preference

**DOI:** 10.64898/2026.09.03.749214

**Authors:** Leslie A. Ramsey, Katherine E. Savell, Rajtarun Madangopal, Caitlyn N. M. Dulan, Fernanda M. Holloman, Samantha S. Lee, Stephanie Oh, M. Flavia Barbano, Marco Venniro, Christopher T. Richie, F. Javier Rubio, Bruce T. Hope

## Abstract

Learned associations between drugs of abuse and drug-associated stimuli are thought to be encoded by physical alterations, termed engrams, within activity-dependent neuronal ensembles. However, the molecular mechanisms that support drug-memory engrams remain largely unknown. The goal of our study was to assess transcriptional alterations as candidate engrams within medial prefrontal cortex (mPFC) ensembles selectively activated by exposure to a cocaine-paired context following cocaine conditioned place preference (CPP). Using the immediate early gene Fos to identify strongly activated neurons, we found that mPFC was strongly activated by exposure to the cocaine-paired context, but not to the unpaired saline control context. We then used a combination of fluorescence activated cell sorting (FACS) and subsequent qPCR to identify gene targets upregulated in the Fos-positive mPFC ensemble neurons relative to the Fos-negative mPFC neurons following context re-exposure. Of the genes examined, the most strongly induced in the Fos-expressing neurons was *Grin2c*, which encodes the NR2C subunit of the NMDA receptor. We confirmed this finding using RNAscope *in situ* hybridization. To assess a causal role for *Grin2c* in cocaine-induced CPP, we used AAV-mediated, neuron-selective expression of *Grin2c* microRNA to knockdown *Grin2c* expression in mPFC neurons, which reduced cocaine CPP and facilitated CPP extinction over repeated tests. Our results indicate that neuronal *Grin2c* expression in the mPFC plays a role in associative learning underlying cocaine CPP. Future studies are needed to determine whether *Grin2c* expression plays a broader role within engrams underlying other forms of learning and memory.

## Introduction

Animals learn to associate the rewarding effects of addictive drugs with cues in the drug-taking environment that can later promote craving and relapse long after the acute effects of the drugs have subsided^1–3^. These memories are encoded within specific patterns of sparsely distributed neurons, called neuronal ensembles, that are activated specifically by the drug-associated cues during learning and memory retrieval^4–6^. The immediate early gene *Fos*^7^ has often been used to label strongly activated neurons during memory retrieval and to demonstrate causal roles for these neuronal ensembles in many learned behaviors, including food^8–10^ and drug reward-based associative learning^11–19^. We and others identified molecular and cellular alterations within these ensemble neurons^20–24^ that are thought to form the long-lasting physical traces called engrams^4^ that encode the food and drug reward memories^5^.

The medial prefrontal cortex (mPFC) plays a key role in drug reward learning in both humans^25–29^ and animal models^30–42^. Neuronal ensembles in mPFC have also been shown to play causal roles in drug-related learned behaviors^18, 19, 43, 44^. While several molecular and electrophysiological alterations have been identified in these ensemble neurons after drug exposure^45–48^, the search for the molecular, cellular, and synaptic substrates of drug reward learning engrams within the mPFC is only just beginning.

Here, we used cocaine conditioned place preference (CPP) to establish a long-lasting cocaine context-associated memory in C57BL/6J mice^49–51^ and assessed gene expression changes in ensemble neurons within the mPFC, with a focus on targets that were known to have effects on neuronal function. We combined Fos immunohistochemistry with fluorescence-activated cell sorting (FACS), and qPCR, to identify transcriptional alterations in mPFC neurons activated during recall of the cocaine-paired context. Amongst other genes, we found *Grin2c* was highly enriched in the activated mPFC neuronal population after context re-exposure. *Grin2c* encodes the NMDA receptor subunit 2c (GluN2C or NR2C)^52^ and has been implicated in a diverse array of functions, ranging from Alzheimer’s disease and cognitive effects of sleep deprivation^52, 53^ to aversion-resistant alcohol intake and intracranial self-stimulation^54, 55^. Our results were unexpected given that Grin2c is only sparsely expressed in the cortex under basal conditions, and furthermore cortical Grin2c expression is primarily restricted to astrocytes rather than neurons^56^. RNAscope *in situ* hybridization confirmed that context exposure selectively increased *Grin2c* expression in Fos-positive neurons, with the highest percentage of Fos-positive neurons that express Grin2c in mice that were exposed to the cocaine-paired context. To assess the functional relevance of neuronal *Grin2c* expression, we employed a viral microRNA-based knockdown strategy targeting mPFC neurons and evaluated the effect of *Grin2c* knockdown on cocaine CPP. We found a reduction in cocaine CPP, suggesting that neuronal *Grin2c* expression plays a causal role in drug-related associative memories, and thus may contribute to the addicted engram.

## Methods and Materials

Subjects. We used a total of 155 male C57BL/6J mice from Jackson laboratories (Bar Harbor, ME, USA), between the ages of 8-10 weeks old at the beginning of our experiments. Mice were maintained in a temperature and humidity-controlled facility on a 12 h reversed light/dark cycle. Mice were separated and individually housed 3-5 days before training and maintained in single-housing throughout the duration of the experiment. Conditioning sessions were conducted in the morning and afternoon during the dark cycle. Food and water were available *ad libitum* throughout the duration of all experiments. All procedures were approved by the NIDA IRP Animal Care and Use Committee (ASP# 23-BNRB-201) and followed the guidelines outlined in the Guide for the Care and Use of Laboratory Animals. We only used male mice because the behavioral study was performed before the implementation of the NIH sex as a biological variable guideline.

### Conditioned Place Preference (CPP) Apparatus

We used a three-chamber apparatus consisting of two equal-sized chambers separated by a smaller middle chamber (59.6 x 17.8 x 15.2 cm). The compartments contained visual and tactile cues to differentiate the chambers. One compartment had black and white striped walls with a grid floor while the other compartment had solid gray walls and a mesh floor. We recorded testing and conditioning sessions using CaptureStar software and analyzed videos using TopView Behavioral Analysis software (CleverSys, Inc. Reston, VA).

### Cocaine CPP

Mice were habituated to the experimenter and weighed daily for a week prior to the first pretest. During the 15 min pretest, mice were allowed to explore all three chambers of the apparatus to measure baseline side preference. We excluded mice that spent >60% of the pretest time in one side of the apparatus, or >30% of the pretest time in the central chamber (2 mice were excluded based on these criteria). During conditioning sessions, mice received a saline injection in one chamber and an injection of cocaine HCl (10 mg/kg, i.p.; NIDA pharmacy) in the other chamber. Injection context and order of injection were counterbalanced between groups. We conditioned mice for 3 days, with one 30 min session in the morning and one 30 min session in the afternoon separated by a 5–6 hour period. Mice were allowed to explore the three chambers during the 15 min test. For experiment 1 (Fig. 1), mice were retested 10 days later, and then re-exposed to the cocaine or saline-paired context 7 days after the retest. Mice were euthanized, then brains were extracted and processed for Fos immunohistochemistry 90 minutes after the beginning of context re-exposure. For experiment 2 (Fig. 2A) and Experiment 3 (Fig. 4A), the CPP training procedure was identical, but mice were re-exposed to the cocaine or saline-paired context 10 days after the test. Then mice were euthanized and brains were extracted 90 min later for either FACS sorting and qPCR, RNAscope *in situ* hybridization. For experiment 4 (Fig. 5A), CPP training procedure was identical, but mice were retested 10 days later and were given a final test 7 days after the retest before they were euthanized to check virus injection placements. For Fig. 5D, mice were exclusively re-exposed to the cocaine-paired side of the chamber.

**Figure 1.**
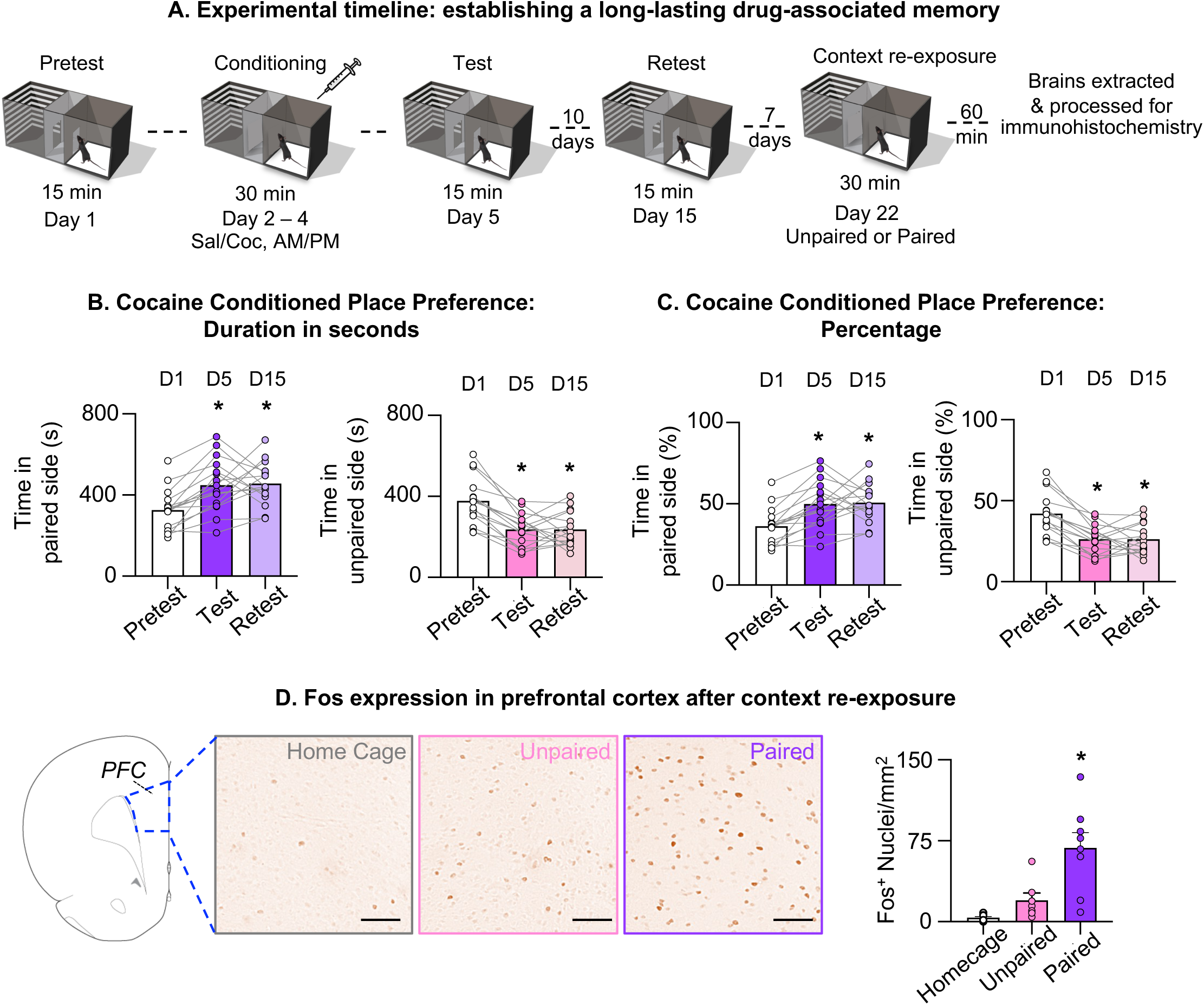
Cocaine conditioned place preference produces a long-lasting drug-associated memory and elevated Fos expression in medial prefrontal cortex (mPFC) upon cocaine-paired context re-exposure. **(A)** Experimental timeline. **(B)** Time (s) spent in the cocaine-paired and saline-paired (unpaired) chambers at pretest, test, and retest. Mice spent significantly more time in the paired side and less time in the unpaired side at test and retest compared with pretest (1-way ANOVA, [F_(1.6,24)_ = 11.78], *p < 0.05 vs. pretest; n=16). **(C)** Same data expressed as percentage of total session time in the paired and unpaired chambers (*p < 0.05 vs. pretest). **(D)** Representative images and quantification of Fos-immunoreactive nuclei in prefrontal cortex (PFC) following exposure to the home cage, unpaired context, or paired context. Paired context re-exposure produced significantly greater Fos⁺ nuclei density than home cage/unpaired conditions (1-way ANOVA, [F_(2,20)_ = 13.56], *p < 0.05; n = 7-8 per group). Data are mean ± SEM.

**Figure 2.**
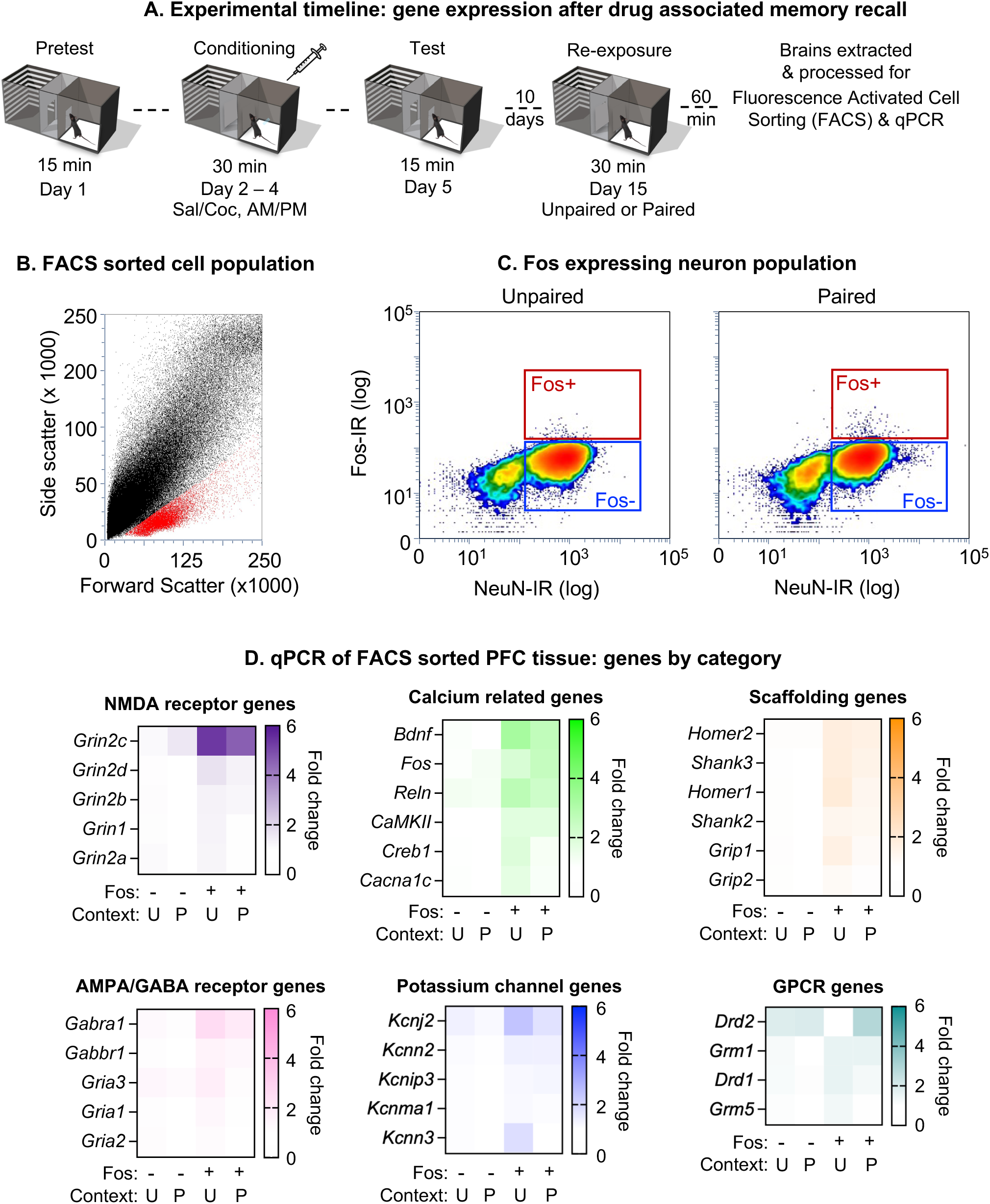
FACS isolation of Fos-expressing mPFC neurons and qPCR analysis following cocaine-paired context re-exposure. **(A)** Experimental timeline. **(B)** Representative FACS gating strategy based on forward and side scatter to isolate single-cell suspensions from PFC tissue. Cell population is shown in red. **(C)** Representative gating of Fos-immunoreactive (Fos-IR) versus NeuN-immunoreactive (NeuN-IR) populations, defining Fos⁺ and Fos⁻ neuronal subpopulations, in mice re-exposed to the unpaired (left) or paired (right) context. **(D)** Heat maps showing fold change in expression (relative to Fos⁻/unpaired) of genes grouped by functional category: genes that encode NMDA receptor subunits, AMPA/GABA receptor subunits, calcium-related genes, genes that encode potassium channels, scaffolding proteins, and GPCRs in Fos⁻ and Fos⁺ populations sorted from unpaired and paired context-exposed PFC. (n = 34 mice, 2 mice pooled per sample; n=9 unpaired samples, n=8 paired samples).

**Figure 3.**
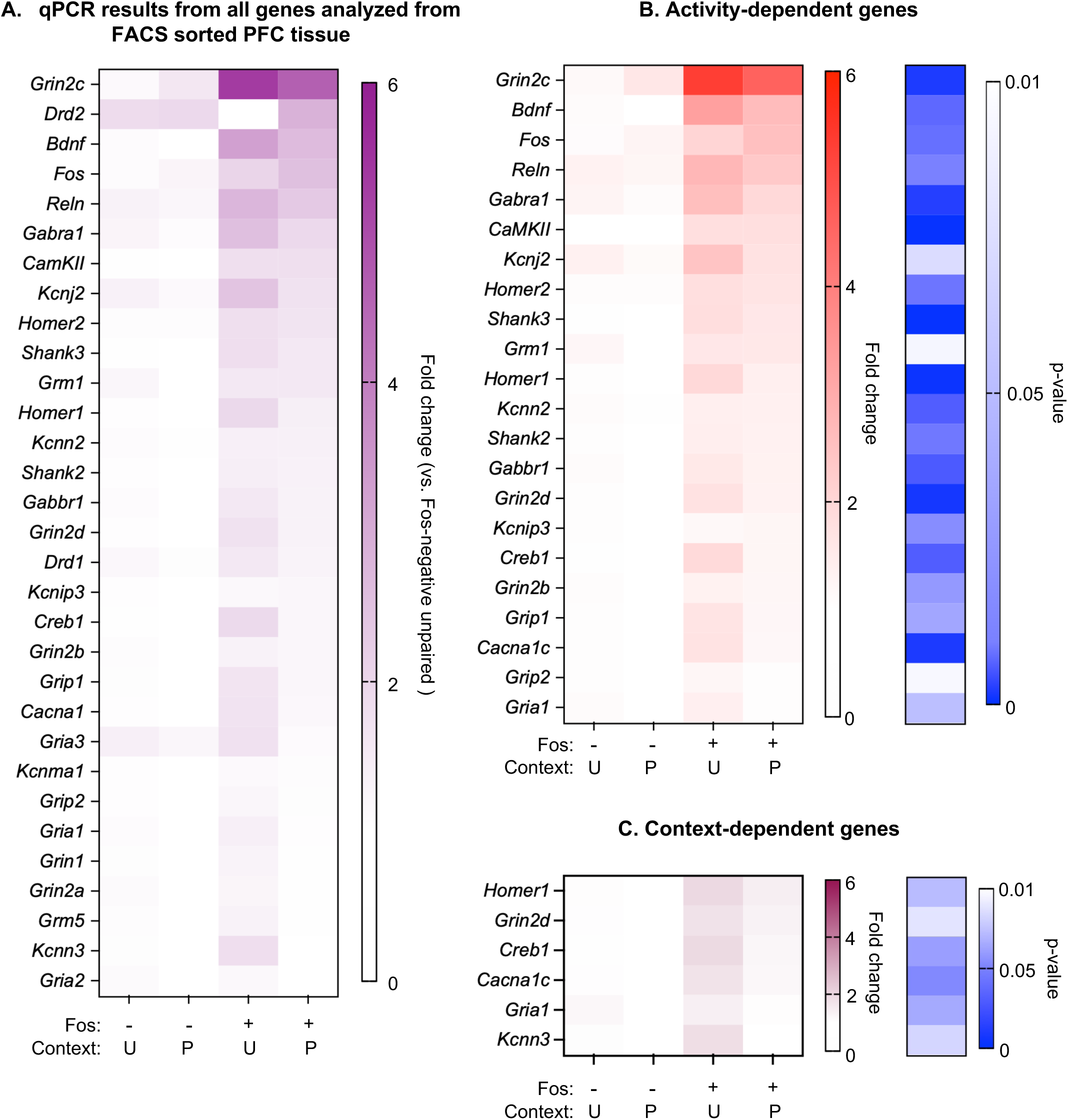
Activity and context-dependent modulation of gene expression after context re-exposure in Fos-positive mPFC neurons. **(A)** Heatmap of fold change (relative to Fos-negative, unpaired condition) for all genes analyzed by qPCR in FACS-sorted PFC tissue from Fos⁻/Fos⁺ neurons following unpaired or paired context re-exposure, ranked by highest to lowest fold change in the Fos+ paired group. **(B)** Genes significantly modulated by activity in the Fos+ population regardless of context (left heatmap), with corresponding p-values (right heatmap). **(C)** Genes significantly modulated by context in the Fos⁺ population, with corresponding p-values (right heatmap). Statistical comparisons were performed using 3-way ANOVA (Significant Gene x Activity interaction, [F_(29,371)_ = 6.01], *p < 0.0001, n=8-9/group) followed by separate 2-way ANOVA for each gene with multiple comparison correction (n = 34 mice, 2 mice pooled per sample; n=9 unpaired samples, n=8 paired samples).

**Figure 4.**
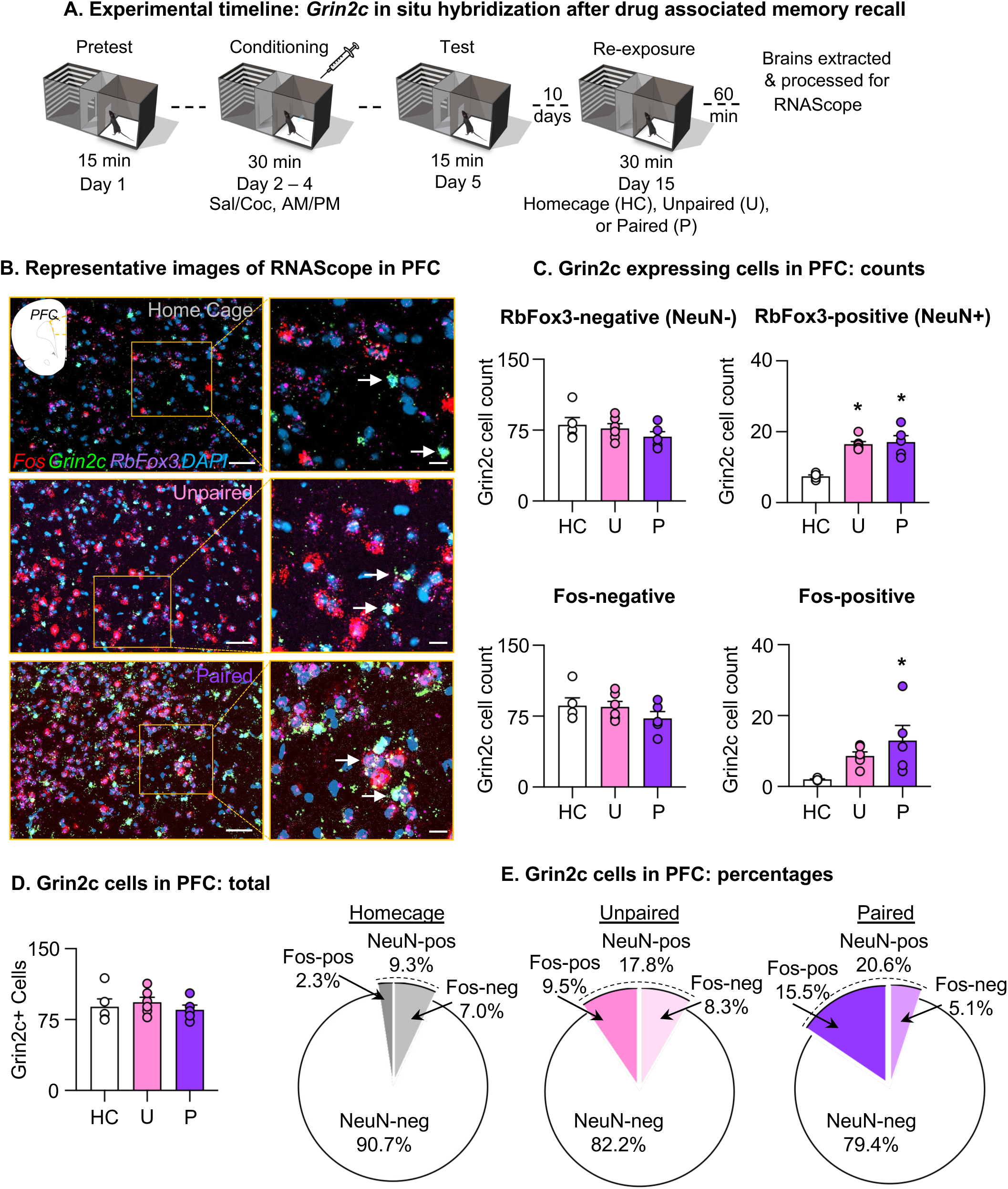
*Grin2c* expression is increased in Fos-positive mPFC neurons following cocaine-paired context re-exposure. **(A)** Experimental timeline. **(B)** Representative confocal images of mPFC sections labeled for *Fos* (red), *Grin2c* (green), *RbFox3/NeuN* (purple), and DAPI (blue) in homecage (HC), unpaired (U), and paired (P) conditions; arrows indicate *Grin2c*⁺ cells. Scale bars, 50 µm (main panel) and 20 µm (inset). **(C)** Quantification of *Grin2c*-expressing cell counts in mPFC, grouped by NeuN status (NeuN⁻, NeuN⁺) and Fos status (Fos⁻, Fos⁺), across HC, U, and P conditions. *Grin2c*⁺ cell counts were significantly increased among NeuN⁺ cells in the U and P groups relative to HC (1-way ANOVA, [F_(2,13)_ = 23.00]; *p < 0.0001; n=5-6/group), and among Fos⁺ cells selectively in the P group (1-way ANOVA, [F_(2,13)_ = 4.90]; *p < 0.05; n=5-6/group). **(D)** Total *Grin2c*⁺ cell counts in mPFC did not differ across HC, U, and P conditions. **(E)** Pie charts depicting the percentage of total *Grin2c*⁺ cells that are NeuN⁻, NeuN⁺/ Fos⁻, and NeuN⁺/Fos⁺ in HC, U, and P conditions, illustrating a shift toward Fos-co-expressing neuronal *Grin2c*⁺ cells with paired-context exposure. Data are mean ± SEM.

**Figure 5.**
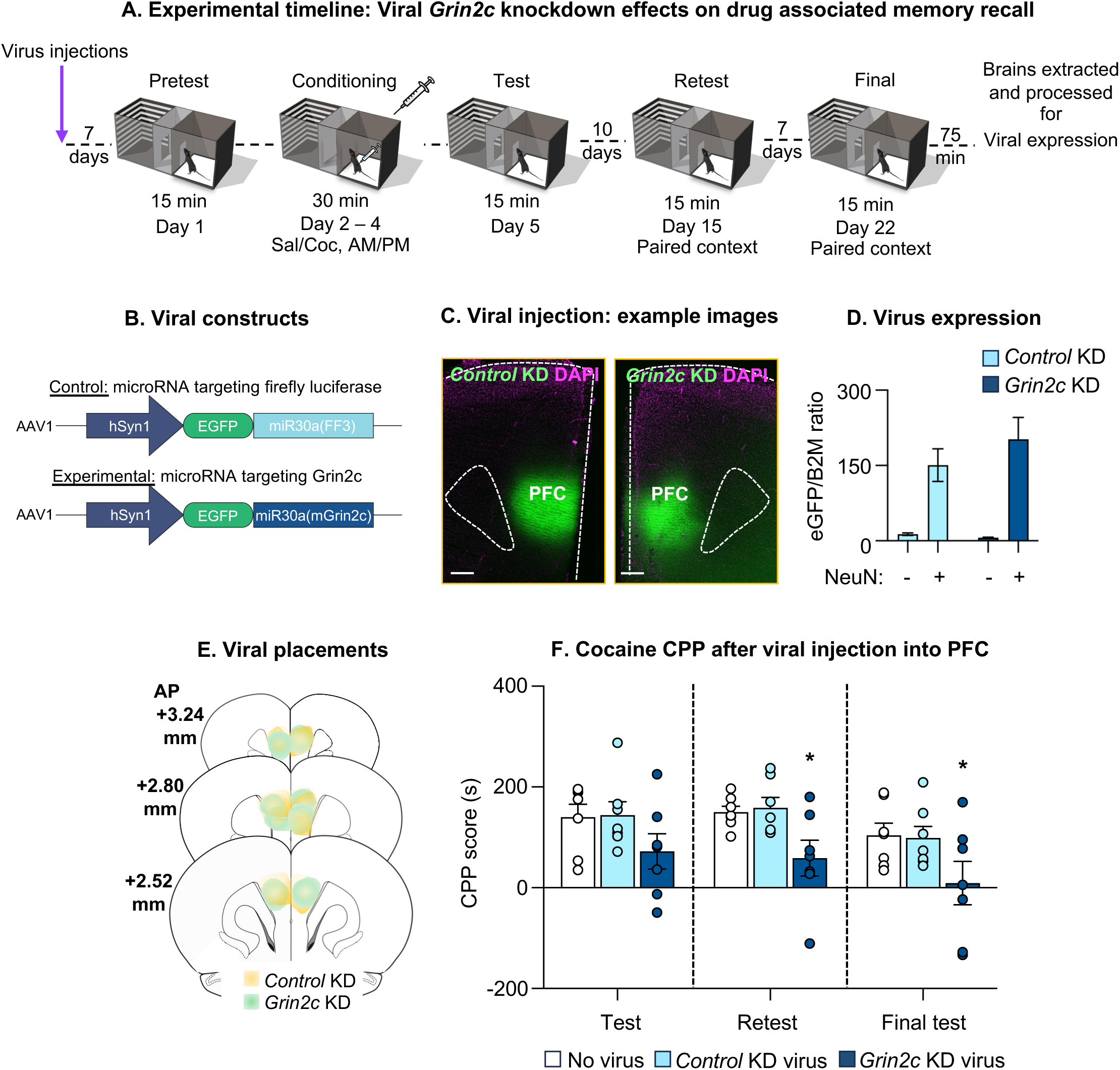
Viral knockdown of *Grin2c* in mPFC neurons attenuates cocaine conditioned place preference. **(A)** Experimental timeline. **(B)** Viral constructs: control virus expresses EGFP with a microRNA targeting firefly luciferase [miR30a(FF3)]; experimental virus expresses EGFP with a microRNA targeting *Grin2c* [miR30a(mGrin2c)], both under the hSyn1 promoter. **(C)** Representative images of mPFC sections showing EGFP expression (green) and DAPI (magenta) for control knockdown (KD) and *Grin2c* KD virus injections. Scale bars, 50 µm. **(D)** Quantification of viral expression (eGFP/B2M ratio) in NeuN⁻ and NeuN⁺ mPFC populations for control KD and *Grin2c* KD groups (2-way ANOVA, main effect of NeuN expression: (2-way ANOVA, [F_(1,13)_ = 38.44], *p < 0.0001; n=7-8/group). **(E)** Viral injection placements within mPFC relative to bregma for control KD (yellow) and *Grin2c* KD (teal) groups. **(F)** Cocaine CPP score (time in paired side minus time in unpaired side, s) at test, retest, and final test for mice receiving no virus, control KD virus, or *Grin2c* KD virus. *Grin2c* knockdown significantly reduced CPP score at retest and final test relative to control groups (2-way ANOVA, [F_(2,18)_ = 4.30], *p < 0.05; n = 7-8/group), indicating impaired persistence of the drug-context memory. Data are mean ± SEM.

### Fos immunohistochemistry

Brains were post-fixed in 4% PFA for 2-3 hours and kept in 30% sucrose at 4°C for 2-3 days. Coronal slices (40 µm) containing mPFC were cut using a cryostat (Leica CM1850). For Fos-labeling experiments, sections were incubated at 37°C for 24 h with a phospho c-Fos rabbit monoclonal antibody (1:1000 dilution, Cat# 5348S, Cell Signaling Technologies, RRID: AB_10557109). Sections were then incubated at 22°C for 2 h in biotinylated goat anti-rabbit secondary antibody (1:600 dilution; Cat# BA-1000, Vector Laboratories, RRID:AB_2313606). Sections were then incubated for 1 h in ABC Elite Kit (Cat# PK-6100, Vector Laboratories). Sections were then reacted in diaminobenzidine for 1-3 minutes. Sections were mounted, air-dried and coverslipped with Vectashield fluorescent mounting medium (Vector Laboratories). We digitally captured bright-field images of immunoreactive (IR) cells in mPFC using an EXi Aqua camera (QImaging) attached to a Zeiss Axioskop 2 microscope at 200× magnification (Carl Zeiss Microscopy) and iVision software for Macintosh v4.0.15 (BioVision). We quantified Fos labeling from four hemi-sections using ImageJ (4 images per mouse) and divided total cell counts by the area quantified to determine mPFC Fos-positive cell density for each subject. Experimenters were blinded to treatment groups during image quantification.

### Fluorescence activated cell sorting (FACS)

Mice were anesthetized with isoflurane and decapitated 30-60 sec later. Brains were removed and tissue was flash frozen in -40C isopentane and then stored at -80C. Three hundred µm sections containing mPFC (AP +1.50 mm to +2.50 mm) were cut on the cryostat (Leica CM 3050S) and tissue punches were taken using a 1.2 mm puncher. We pooled 2 mouse mPFCs per tissue sample. Tissue was processed to obtain dissociated cells as previously described^57^. Briefly, tissue was finely minced with razor blades and placed on ice in a tube containing 1 ml of Hibernate A (catalog #HA-lf, Brain Bits) and enzymatically digested for 30 min at 4°C with 1 ml of Accutase (catalog #SCR005, Millipore). Each tissue sample was triturated three times in series using fire-polished glass pipettes with successively smaller diameters (1.3, 0.8, and 0.4 mm). Samples were fixed and permeabilized in final 50% cold ethanol for 15 min. Cells were centrifuged for 4 min at 1000 × *g* and resuspended in 1 ml of cold PBS. Then fixed cells were filtered through 100 and 40 μm cell strainers (Falcon, BD Biosciences). Cell suspensions were split into two tubes for each sample. One tube (700 μl) was incubated with the primary and secondary antibodies for sorting cells. The second tube (remainder of each cell suspension after filtering, ∼250 μl) was used to set fluorescence thresholds for positive NeuN-IR and Fos-IR gates as well as to detect total cell populations with DAPI.

We used phycoerythrin (PE)-conjugated mouse monoclonal antibody against NeuN (1:500 dilution; catalog #FCMAB317PE, clone A60, Millipore, RRID: AB_11212465) and Alexa Fluor 647-conjugated rabbit monoclonal phospho-c-Fos antibody (1:500 dilution; catalog #8677, Cell Signaling Technologies, RRID:AB_11178518) for FACS. The cell suspension was incubated with primary antibodies for 30 min at 4°C. Cells were washed twice with cold PBS followed by centrifugation for 3 min at 425 × *g*. The pellet was resuspended in 0.5 ml of cold PBS and samples were sorted using a FACSAria Fusion SORP flow cytometer (BD Biosciences). As we previously reported, cells containing neurons can be identified by their unique forward scattering (FSC) and side scattering (SSC) properties^57^.

A control sample stained with the nuclear counterstain DAPI was used to define the ‘Cells’ gate from the total events and cellular debris before sorting the main samples. Between 80-95% of the events in the cells and neurons gates, respectively, were DAPI-positive events (cells with nuclei). A gating strategy based on FSC width and height was used to select only single cells before gating for Fos-positive and Fos-negative neurons. We sorted neurons according to phycoerythrin (PE; NeuN-immunopositive) and Alexa Fluor 647 (Fos-immunopositive) fluorescence signals. A maximum of 5000 NeuN-positive and Fos-negative events (Fos-negative neuron population) and all NeuN-positive and Fos-positive events (Fos-positive neuron population) were sorted. For the data shown in Fig. 5D, we used the same tissue processing (no ethanol fixation) and staining steps, and sorted all NeuN-positive events and NeuN-negative events.

### RNA isolation and cDNA synthesis from FACS-isolated neurons

We used FACS with the multiplex preamplification strategy as described previously to analyze gene expression in mouse mPFC neurons^58^. Briefly, sorted cells were collected directly from FACS into 50 μl of the extraction buffer from PicoPure RNA isolation kit (Arcturus Bioscience) and lysed by pipetting up and down 10 times followed by incubation at 42°C for 30 min. The suspension was centrifuged at 1000 × *g* at 4°C for 2 min and the supernatant was collected for RNA isolation. Column filtration, washing, and elution steps of RNA from the columns are described in section C of the PicoPure RNA isolation protocol. This protocol reliably produces high-quality RNA for gene expression analysis^20, 57, 59–61^. Single-strand cDNAs were synthesized with the Superscript III first strand cDNA synthesis kit according to the manufacturer’s protocol (Invitrogen, Life Technologies). For data shown in Figure 5D, RNA was isolated from the sorted populations using a QIAGEN EZ2 Connect MDx robot and QIAGEN RNA/miRNA Tissue/Cells prep kits.

cDNA preamplification was performed using the TaqMan PreAmp Master Mix Kit according to the manufacturer’s general workflow (Applied Biosystems/Life Technologies). A pooled primer/probe solution was prepared by diluting each 20X TaqMan Gene Expression Assay 1:100, resulting in a 0.2X concentration for each TaqMan assay in the pooled solution. Custom-designed primer sets were included in the pooled solution at 80 nM each (Supplementary Table 2). For each preamplification reaction, 7.5 μl of cDNA was combined with 7.5 μl of the pooled primer/probe solution and 15 μl of 2X TaqMan PreAmp Master Mix, for a final reaction volume of 30 μl. Thus, the final concentrations in the preamplification reaction were 0.05X for each TaqMan Gene Expression Assay and 20 nM for each custom primer set. cDNAs were preamplified in an Applied Biosystems 9700 Thermal Cycler using the following cycling conditions: an initial hold at 95°C for 10 min, followed by 14 cycles of denaturation at 90°C for 15 s and annealing/extension at 60°C for 4 min. The preamplification PCR products were immediately diluted fivefold with molecular biology-grade water (5 Prime) and either stored at −20°C or processed immediately for qPCR.

For data shown in Figures 2 and 3, duplex qPCR assays were performed in technical duplicate using a FAM-labeled probe for each target gene and a VIC-labeled probe for the endogenous control gene, *RbFox3*, which encodes the NeuN protein, together with TaqMan Advanced Fast PCR Master Mix (Life Technologies). *RbFox3* was selected as the endogenous control because it was relatively abundant, not altered by the CPP procedure, and did not differ between Fos-positive and Fos-negative neurons (data not shown). Primer Express 3.0 (Applied Biosystems) was used to design primers and probes for preamplification and qPCR. Primers and probes were selected to amplify across exon–exon junctions for each target gene. qPCR was performed on a 7500 Fast TaqMan instrument using the following cycling conditions: an initial hold at 95°C for 20 s, followed by 40 cycles of denaturation at 95°C for 3 s and annealing/extension at 60°C for 30 s. Relative gene expression was calculated from Ct values using the ΔCt method, where ΔCt = Ct target gene − Ct RbFox3. For each sample, the mean ΔCt value of Fos-negative neurons from mice tested in the Unpaired context was subtracted from the sample ΔCt value to generate a ΔΔCt value. Fold change relative to the Unpaired Fos-negative group was then calculated as 2^−ΔΔCt^62^. For data shown in Figure 5D, quantification of target template (EGFP) and a reference housekeeping gene (mB2M) was performed using multiplex probe assays with a QIAGEN QIAcuity digital PCR machine. The target template concentration for each reaction was normalized to the housekeeping gene, then the technical replicates were averaged together.

### RNAscope ISH assay

We anesthetized mice and then extracted the whole brain, which was frozen for 20 s in 100 ml −50°C isopentane within 2–3 min of decapitation, wrapped in labeled aluminum foil sealed in a zippered plastic bag, stored at −80°C, and cryosectioned for ISH. Brains were equilibrated to −20°C in a cryostat (CM 3050S) for 2 h and 20 μm coronal sections containing mPFC (AP +2.2 to +1.8 mm) were cut and thaw-mounted directly onto Super Frost Plus slides (Fisher). These slides were left at −20°C for 10 min and transferred to −80°C until ISH processing.

We manually performed RNA ISH for *Fos*, *Grin2c*, and *RbFox3* mRNAs according to the *User Manual for Fresh Frozen Tissue using RNAscope Multiplex Fluorescent Reagent Kit* (Advanced Cell Diagnostics). Briefly, the −80°C brain slides were transferred to slide racks and fixed by immersion in 10% neutral buffered formalin (Fisher Scientific) for 20 min at 4°C. Slides were rinsed two times in PBS and dehydrated two times each in 50 and 70% ethanol and twice in 100% ethanol. Slides were transferred to a new 100% ethanol container and kept at −20°C overnight. The slides were dried at room temperature (22°C) for 10 min and a hydrophobic pen (ImmEdger Hydrophobic Barrier Pen, Vector Laboratories) was used to make a physical barrier surrounding the brain sections to contain RNAscope assay solution. We used the HybEZ Hybridization System from Advanced Cell Diagnostics. The protease solution (Pretreatment 4 solution) was incubated with sections at room temperature for 20 min. After washing off the protease solution, 1× target probes for specific RNAs (*Fos*, *Grin2c*, and/or *RbFox3*) were applied to the brain sections and incubated at 40°C for 2 h in the HybEZ oven. Each RNAscope target probe contained a mixture of 20 ZZ oligonucleotide probes that bound to the target RNA. These probes were as follows: *Grin2c*-C1 probe (accession number <u>NM_010350.2</u>, target nucleotide region: 1332-2367); *Fos*-C2 probe (accession number <u>NM_010234.2</u>, target nucleotide region: 407-1427); *RbFox3*-C3 probe (accession number <u>NM_001039167.1</u>, target nucleotide region: 1827-3068).

Sections were then incubated with preamplifier and amplifier probes by applying AMP1 (40°C for 30 min), AMP2 (40°C for 15 min), and AMP3 (40°C for 30 min). Sections were then incubated with the fluorescently labeled probes by selecting a specific combination of colors associated with each channel: green (Alexa 488 nm), orange (Alexa 550 nm), and far-red (Alexa 647 nm). We used AMP4 AltB to detect triplex *RbFox3*, *Fos*, and *Grin2c* RNAs in far-red, orange, and green respectively. Finally, we incubated the sections for 20 s with DAPI to stain nuclei (blue). Between each step we washed two times with 1× wash buffer supplied with the kit. The negative control sections received RNase treatment before performing the RNAscope assay; after the fixation and protease digestion, we incubated sections with 5 mg/ml RNase A (Qiagen) for 30 min at 40°C. The slides were washed three times with distilled water and processed with target probe hybridization, followed by the steps described above. We captured fluorescent images of labeled cells in mPFC using a Rolera EM-C^2^ camera (QImaging) attached to a Nikon Eclipse E800 at 200× magnification and iVision software for Macintosh v4.0.15 (BioVision). We quantified mRNA colabeling from two hemi-sections using ImageJ (2 images per mouse) in a blinded manner.

### miRNA AAV vector genome plasmid construction

The short hairpin RNA (shRNA) targeting mGrin2C described by Seif et al (2013)^54^ was adapted into a microRNA cassette using the conversion process previously described^63^. The mGrin2C microRNA cassette was then used to replace the previous KASH domain in pOTTC1730 (pAAV SYN1 EGFP-KASH) using ligation-independent cloning to produce the plasmid pOTTC1984 encoding the genome for the Grin2c KD virus: pAAV-hSyn1-EGFP-miR-30a(mGrin2c). A non-specific control miRNA targeting firefly luciferase (FF3) was amplified from pOTTC1618 (pAAV SYN1 NUC-EYFP mIR-30 FF3) ^64^ and similarly used as an insert to create pOTTC1985 encoding the genome for the Control FF3 KD virus: pAAV-hSyn1-EGFP-miR-30a(FF3). Plasmids will be deposited with Addgene.

### AAV packaging and purification

Each viral vector was packaged as previously described^65^ with minor modifications. HEK293 cells were transfected using calcium phosphate precipitation to deliver plasmids encoding the adenovirus helper genes and the adeno-associated replicase and capsid genes for AAV serotype 1 (pHelper and pXR1, respectively^66^, and a plasmid encoding the vector genome). Cell lysates were collected 40 hours post-transfection, treated with SAN HQ nuclease (Articzymes), and passed through an AVB columns (GE Healthcare, Silver Spring, MD) using a fast protein liquid chromatography (FPLC) machine (AKTA). Purified viral particles were eluted from the column with sodium citrate (pH ∼3.0) and titered using digital PCR (QIAGEN QIAcuity) using a probe-based assay that recognizes the WPRE sequence.

### Intracranial Virus Surgery

We anesthetized mice with isoflurane (4-5% induction, 1-3% maintenance) prior to injecting AAVs bilaterally into the mPFC using the following stereotaxic coordinates from bregma: antero-posterior (AP), 2.0 mm; medio-lateral (ML), +/-0.8 mm; dorso-ventral (DV), -2.2 mm; 10° angle. We used Nanofil syringes (10 uL syringes with 33G injector needles, World Precision Instruments) attached to an ultramicropump (UMP3, World Precision Instruments) and controller (Micro4, World Precision Instruments) to deliver 300 nL of the Grin2c KD virus or Control FF3 KD virus at a rate of 100 nL/min. Following surgery, we injected ketoprofen (2.5 mg/kg, s.c.; Covetrus) daily for 3 days following surgery to relieve pain and decrease inflammation. We allowed mice to recover from surgery for at least 5 days prior to initial behavioral training.

### Experimental designs

Experiment 1: Effect of cocaine reward memory recall on Fos expression in the mPFC. Mice were trained for cocaine CPP as described above and tested for side preference 1 day (test) and 10 days (retest) after training. Mice were re-exposed to the cocaine or saline-paired context 7 days after the retest and then processed for Fos immunohistochemistry 90 minutes after the beginning of context re-exposure. We included an additional group of mice that were taken directly from the home cage as a control group to determine baseline levels of Fos expression in mPFC.

Experiment 2: FACS sorting of Fos-expressing neurons and gene expression analysis after cocaine reward memory recall. CPP training was identical to Experiment 1, but mice were re-exposed to the cocaine or saline-paired context 10 days after the CPP test, and then brains were extracted 90 min after the start of the test session for FACS sorting and subsequent qPCR on sorted samples as described above.

Experiment 3: *Grin2c* RNAscope *in situ* hybridization after cocaine memory recall. CPP training was identical to Experiment 1, but mice were re-exposed to the cocaine or saline-paired context 10 days after the CPP test. A separate group went through identical CPP training but was left in the home cage on test day. Brains were extracted 90 min after the start of the session for RNAscope *in situ* hybridization. We limited our RNAscope image quantification to cells expressing *Grin2c* mRNA.

Experiment 4: Viral Grin2c knockdown effects on cocaine memory recall. We injected the AAV into mPFC as described above at least 5 days before CPP training. CPP training was identical to Experiment 1, but mice were retested 10 days later and then given a final test 7 days after the retest before euthanizing them to check virus injection placements (Fig. 5C).

Experiment 5: Validation of Grin2c knockdown virus expression in neurons. CPP training was identical to Experiment 1, but mice were re-exposed to the cocaine-paired context 10 days after the CPP test. Brains were extracted 90 min after the start of the session for FACS sorting and subsequent qPCR on sorted samples.

### Statistical analysis

We analyzed the behavioral, immunohistochemical, FACS sorted qPCR, RNAscope ISH, and viral knockdown data by one-way or two-way ANOVAs using Prism (Graphpad Software). For FACS-sorted samples, we used the between-subjects factor of Context (Unpaired, Paired) and the within-subjects factor of neuron type (Fos-negative, Fos-positive). For RNAscope ISH, we used the between-subjects factor of Context (Homecage, Unpaired, Paired) and the within-subjects factor of Neuron Type (*Fos*-negative, *Fos*-positive). For viral knockdown we used the between-subjects factor of Virus (No virus, Control KD virus, Grin2c KD virus) and the within-subjects factor of CPP Test (Test, Retest, Final test). Sidak multiple comparison’s test or Fisher’s PLSD was used for *post hoc* analyses when prior ANOVAs indicated significant main or interaction effects (*p* < 0.05). The detailed results of all statistical analyses are shown in Supplemental Table 1.

## Results

### Establishing a long-lasting drug associated memory using cocaine conditioned place preference

In Experiment 1, we used a cocaine CPP procedure to test if male C57BL/6J mice maintain a long lasting preference for a cocaine-paired context (Fig. 1A). We measured the time spent in the cocaine-paired context (Paired) relative to the saline-paired context (Unpaired) during a 15-minute test session that took place one day after completion of conditioning sessions (Test) and then retested the mice 10 days later (Retest). Mice spent more time in the Paired context (F_1.6,24_= 11.78, p< 0.001; Fig. 1B left), and less time in the Unpaired context (F_1.7,25_= 23.97, p< 0.0001; Fig. 1B right) during Test and Retest as compared to the Pretest, indicating they formed a persistent preference for the cocaine-paired side that lasted 10 days. The same data are expressed as a percentage of test time in Fig. 1C. Seven days after Retesting, mice were re-exposed for 30 min to either the Paired or Unpaired context and euthanized 60 minutes later. We quantified Fos-positive nuclei within the medial prefrontal cortex (mPFC) following re-exposure to either context (Fig. 1D) and compared this with a control group that remained in the Homecage and were immediately euthanized to establish baseline levels of Fos expression. Mice re-exposed to the Paired context had increased mPFC Fos expression compared to the other two groups (F _2,20_= 13.56, p<0.001; Sidak’s posthoc test: Homecage vs. Paired: p=0.0002, Unpaired vs. Paired: p=0.005).

### FACS-sorting and qPCR gene expression analysis reveals *Grin2c* upregulation in Fos-positive PFC neurons

In Experiment 2, we trained mice using an identical CPP procedure in which mice demonstrated a similar magnitude of CPP as in Experiment 1 (Fig. S1). Ten days after the CPP Test, mice were re-exposed to either the Paired or Unpaired context (Fig. 2A), and brains were extracted 90 min from the start of the exposure. We used fluorescence-activated cell sorting (FACS) to isolate Fos-positive and Fos-negative neurons (NeuN-positive population, neuronal marker) (Fig. 2B-C) and then qPCR to examine gene expression in the Fos-positive vs. Fos-negative neuronal populations. We selected a list of genes to analyze based on their functional roles in neuronal activity modulation, synaptic plasticity, or drug-based associative learning. We grouped genes into six functional categories: NMDA receptor genes, calcium-related genes, scaffolding genes, AMPA/GABA receptor genes, potassium channel genes, and GPCR genes (Fig. 2D). Gene expression data are presented in terms of fold change (ΔΔCt) relative to the Fos-negative Unpaired group and are aligned in descending order from highest to lowest fold change in the Fos-positive Paired group for each gene category (Fig. 2D) and all genes (Fig. 3). We analyzed gene expression data using a 3-way ANOVA which included between-subjects factors of Context (Paired, Unpaired) and Gene (all listed genes), and within-subjects factor of Activity (Fos-positive, Fos-negative) (Fig. 3A). We identified a significant Gene x Activity interaction (F_(29,371)_ = 6.01, p<0.0001) as well as main effects of Gene (F_(29,448)_ = 14.86, p<0.0001) and Activity (F_(1,371)_ = 86.12, p<0.0001) separately. We also analyzed each gene separately using a 2-way ANOVA with between subjects factor of Context and within subjects factor of Activity. Because our analysis involved screening multiple gene targets, we corrected 2-way ANOVA p-values for multiple comparisons using Benjamini and Hochberg’s false discovery rate (FDR) correction method and set alpha (significance) level at 0.1, two-tailed. Effects meeting this threshold were interpreted as exploratory findings. We found that *Grin2c*, the gene that encodes the NR2C subunit of the NMDA receptor, had the highest fold change in the Fos-positive Paired group relative to all other analyzed genes (Fig. 3A). We did not identify significant interactions between Context and Activity in any of the genes analyzed. However, we identified 22 genes that were modulated by Activity (Fig. 3B), and 6 genes that were modulated by Context (Fig. 3C), which are presented in separate heatmaps with rows ordered in descending order from highest to lowest fold change in the Fos-positive Paired group. *Grin2c* had the highest fold change among the genes modulated by activity as well. In the majority of the genes assessed, there were higher fold changes in the Fos+ Unpaired group than in the Fos-positive Paired group (Fig. 3A). The only exceptions to this rule were the *Drd2* and *Fos* genes.

### RNAscope *in situ* hybridization reveals increased *Grin2c* expression in Fos-positive neurons after drug associated memory recall

Under baseline conditions, *Grin2c* expression is typically undetectable or very low in prefrontal cortical neurons, thus we used RNAscope *in situ* hybridization to independently confirm *Grin2c* enrichment in Fos-positive neurons. In Experiment 3, we trained mice using an identical CPP procedure. Mice demonstrated similar place preference levels relative to Experiments 1 and 2 (Fig. S2). Ten days after the CPP test, mice were re-exposed to the Paired or Unpaired context. A control group of mice remained in the Homecage to determine if baseline *Grin2c* expression levels were similar to what has previously been reported. Following context re-exposure, brains were removed (90 min. after start of the exposure) and processed for RNAScope *in situ* hybridization (Fig. 4A) using highly specific probes to detect expression of *Fos*, *Grin2c*, and *RbFox3 (*the gene that encodes the neuronal marker, NeuN). We detected DAPI-labeled cells expressing *Grin2c* in the Homecage group, but detected very low levels of overlap between *Fos*, *NeuN*, and *Grin2c* in the Homecage group. However, after context re-exposure, we detected triple labeled cells that expressed *Grin2c*, *Fos*, and *NeuN* (Fig. 4B). We found that there were similar numbers of *Grin2c*-expressing cells in the NeuN-negative group (non-neuronal cell types; F_(2,13)_ = 1.07, p=0.37), but more *Grin2c*-expressing cells in the NeuN-positive group (neuronal cell type; F_(2,13)_ = 23.00, p<0.0001) after exposure to either the Paired or Unpaired context relative to the Homecage control group (Fig. 4C). However, there were more *Grin2c* cells in the Fos-positive neurons, but not in the Fos-negative neurons, after exposure to the Paired context (F_(2,13)_ = 4.90, p<0.03), indicating there are more active neurons expressing *Grin2c* after cocaine-paired context re-exposure. The total number of *Grin2c* cells remained similar between groups regardless of context exposure (Fig. 4D). The same data are expressed in terms of percentage in the charts shown in Fig. 4E. Fifteen and a half percent of *Grin2c*-expressing cells are Fos-positive neurons in the Paired context re-exposure group whereas only to 2.3% of *Grin2c*-expressing cells are Fos-positive neurons in the Homecage control group. Our results confirm the presence of Grin2c expression in Fos-positive neurons after context re-exposure that we first identified with FACS and qPCR (Figs. 2 & 3)

### *Grin2c* knockdown in the PFC attenuates cocaine conditioned place preference

Having established via two separate methods that *Grin2c* expression increases in mPFC of mice re-exposed to the cocaine-paired context, we sought to determine whether *Grin2c* expression is necessary for cocaine CPP (Fig. 5A). To do this, we designed a human Synapsin (hSyn1) promoter-driven construct that expresses a microRNA targeting *Grin2c* to knockdown *Grin2c* expression primarily in neurons. An hSyn1-promoter driven construct expressing a microRNA targeting a control gene, Firefly luciferase (FF3), served as our control (Fig. 5B). Mice received bilateral mPFC injections of either Grin2c knockdown virus or the control FF3 knockdown virus prior to CPP training. Using confocal microscopy, we detected eGFP signal in mPFC tissue taken from mice that received either virus (Fig. 5C). We quantified viral expression by taking mPFC tissue punches from mice that received either the control or Grin2c knockdown virus. We used FACS to sort the samples into NeuN-positive (neuronal) and NeuN-negative (non-neuronal) cell populations. We ran qPCR on the sorted samples and quantified eGFP expression relative to the housekeeping gene *B2M,* which encodes the protein ß-2-microglobulin. Our 2-way ANOVA analysis included the between-subjects factor of virus (Control, *Grin2c* knockdown) and the within-subject factor of cell type (NeuN-positive or NeuN-negative), and identified a main effect of cell type (F _1,13_= 38.44, p< 0.0001). We detected very low levels of eGFP in the NeuN-negative cell population regardless of virus type, suggesting that viral expression was restricted to the neuronal, or NeuN-positive cell population for both control and *Grin2c* knockdown viruses (Fig. 5D). After CPP training, we tested place preference at three different time points. Our two-way ANOVA analysis included between-subjects factor of Virus (No Virus, Control, *Grin2c* knockdown) and within subject factor of Test session (Day 5, Day 15, Day 22). We observed a significant main effect of Test session (F _2,36_= 6.72, p= 0.01) and Virus (F _2,18_= 4.30, p=0.03). Posthoc analysis revealed that mice expressing the *Grin2c* knockdown virus in mPFC had significantly lower CPP scores than Control or No Virus groups on the Day 15 and Day 22 test sessions, with a trend toward lower CPP scores on Day 5. These findings suggest that neuronal *Grin2c* expression is necessary for long term drug-associated memory recall.

## Discussion

We used conditioned place preference (CPP) to establish a robust learned association between an environmental context and the rewarding effects of cocaine^49–51^, followed by cocaine-paired context re-exposure to prompt mice to recall the context-cocaine association. We observed that cocaine CPP persists for at least ten days after the first CPP test, in line with reports indicating that cocaine CPP can persist up to a month after the initial test^67, 68^. Exposure to the cocaine-paired context, but not the saline-paired context (unpaired), increased Fos expression in the medial prefrontal cortex (mPFC). In a separate experiment, we used FACS to separate Fos-positive and Fos-negative neurons from mPFC of mice exposed to the cocaine-paired and unpaired contexts and used qPCR to measure expression of 31 genes chosen for their effects on neuronal activity, excitability, or plasticity. Twenty-two of the 31 genes analyzed increased in Fos-positive, but not Fos-negative neurons, following exposure to both the cocaine-paired and unpaired contexts. The gene with the highest fold increase after exposure to either context was *Grin2c*, which encodes the NMDA receptor subunit NR2C. We verified that *Grin2c* expression increases after context re-exposure using RNAscope *in situ* hybridization. *Grin2c* expression was increased in neurons following exposure to either the paired or unpaired contexts, but this time only in Fos-positive neurons after cocaine-paired context re-exposure. To determine whether *Grin2c* expression in mPFC neurons contributes to cocaine CPP, we injected the mPFC with AAVs expressing a *Grin2c*-specific miRNA under a neuron-selective promoter to knock down *Grin2c* expression specifically in neurons. When injected prior to CPP training, *Grin2c* knockdown decreased CPP during the retest and final test, indicating *Grin2c* expression in Fos-expressing mPFC neurons plays a causal role in cocaine CPP.

### A role for mPFC in drug-associated learning and memory

Extensive preclinical and clinical literature implicates the mPFC in drug-based associative learning (for clinical reviews, see^69, 70^; here we focus on preclinical models). The mPFC is active during cocaine self-administration and cocaine- or cue-driven reinstatement to drug seeking^33, 71^, and its inactivation reduces cue-induced reinstatement to cocaine seeking^72, 73^. The mPFC also plays a causal role in cocaine CPP^74–76^. Other studies have observed neuronal activity or plasticity within mPFC cells after cocaine CPP^45^, consistent with our finding that exposure to the cocaine-paired, but not unpaired, context increased Fos expression in mPFC neurons, indicating strong neural activation during recall of the cocaine CPP memory. In previous studies, we and others have found that Fos-expressing neurons in mPFC^19, 43, 77, 78^ and other brain regions^13, 15–17, 79–81^ act together as neuronal ensembles to mediate recall of drug-associated memories.

#### Gene expression changes in Fos-positive neurons after cocaine CPP and context re-exposure

Long-lasting molecular and cellular alterations within Fos+ neurons are thought to represent candidate functional/physical substrates, or engrams, underlying long-term memory. To identify such candidates, we assessed a panel of genes implicated in synaptic plasticity, calcium-dependent activity, neuronal excitability, and neurotransmission, several of which are known to alter synaptic transmission during learning. Many of these genes were upregulated in Fos+ neurons from both the cocaine-paired and unpaired groups. Interestingly, the fold-increase for most of these genes was greater in the unpaired group than in the paired group, despite the relatively low number of Fos-immunolabeled neurons in the unpaired group. Of the genes assessed, 22 were modulated by activity (the top six most induced are *Grin2c*, *Bdnf*, *Fos*, *Reln*, *Gabra1*, *CamKII*) while only 6 were modulated by context (*Homer1*, *Grin2d*, *Creb1*, *Cacna1c*, *Gria1*, *Kcnn3*) after cocaine or saline-paired context re-exposure. Genes upregulated in Fos+ neurons specifically after exposure to the cocaine-paired context represent promising candidate targets warranting further investigation into their causal role in the long-term encoding of cocaine memories.

### Expression of Grin2c in Fos-positive neurons after cocaine CPP and context re-exposure

We chose to focus on the highly induced *Grin2c* gene for further investigation as a candidate engram in this study. *Grin2c* has not been extensively studied in the context of preclinical drug reward learning models, and the large fold induction of *Grin2c* in Fos-positive neurons was somewhat surprising. In the Allen Brain Cell Atlas, *Grin2c* is expressed at low levels in mouse cortex and is most prevalent in the astrocytic cell population rather than the neuronal population^56^; however, gene expression in these databases is assessed in largely non-activated neurons from animals taken directly from their non-arousing home cages. This suggests that neuronal activity or plasticity triggered by re-exposure to the drug-paired context may induce *Grin2c* expression in mPFC neurons. We sorted our samples by activity status (Fos+ vs Fos-) and by cell type (Neun+ vs. Neun-), so the elevated *Grin2c* expression we observed with FACS sorting and subsequent qPCR occurred primarily in active neurons. We used RNAscope *in situ* hybridization with probes targeting *Fos*, *RbFox3,* and *Grin2c* to confirm our finding that Grin2c expression increased in neurons after context re-exposure. Using this method, we identified more *Grin2c* expressing neurons after exposure to either the paired or unpaired context, and we found the highest number of *Grin2c* expressing neurons in the Fos-positive paired group. Interestingly, there was no change in the overall number of cells expressing *Grin2c*, just a shift in the ratio of cell types expressing *Grin2c*. It is likely that the overall numbers of *Grin2c* expressing cells did not change because *Grin2c* expressing neurons still represent a relatively small proportion relative to the number expressed in astrocytes.

### A causal role for neuronal Grin2c expression in cocaine CPP

To determine whether *Grin2c* expression in mPFC is necessary for cocaine CPP, we knocked down *Grin2c* expression specifically in mPFC neurons using a *Grin2c* KD virus with a neuron-specific promoter driving co-expression of *Grin2c*-specific microRNA and GFP as a fluorescent marker. We found CPP scores were lower in mice that received injections of the *Grin2c* KD virus into mPFC compared to mice that received either the control virus or no virus on the Day 15 and Day 22 CPP tests. While the initial test on Day 5 test showed lower CPP scores in the *Grin2c* KD mice, this did not reach statistical significance (p = 0.08). To confirm that the behavioral effect was due to *Grin2* KD specifically in neurons, we FACS sorted neuronal and non-neuronal cells from mPFC injected with the *Grin2c* KD virus that co-expresses GFP and found nearly all co-pressed GFP was in the neuronal population. Thus, we concluded that the *Grin2c* knockdown effects on CPP were due to neuron-specific *Grin2c* knockdown. It should be noted that we were not able to detect a reduction in *Grin2c* expression in mPFC neurons in the *Grin2c* KD mice because the proportion of neuronal *Grin2c* was below the detection threshold of our assays and likely confounded by small numbers of high *Grin2c*-expressing astrocytes in the neuron populations. Seif et al previously demonstrated that shRNA-mediated knockdown of *Grin2c* in the NAc core led to a reduction in aversion-resistant alcohol intake in rats^54^. However, since the knockdown in this study was not cell type-specific, it is not known whether the effect they observed on alcohol intake in rats was due to neuronal or astrocytic *Grin2c* knockdown.

### Grin2c and the nature of its role in drug-associated learning and memory

Several questions arise in light of our results examining activity-dependent gene expression changes and neuronal *Grin2c* knockdown effects. *Grin2c* expression was induced in the Fos-expressing population of mPFC neurons, but it is unclear whether *Grin2c* expression is acutely induced in response to strong neuronal activation, or if it is driven by associative learning specifically, likely as a result of long-term synaptic plasticity. It is also unclear if the effects of *Grin2c* expression are specific to cocaine CPP acquisition (the formation of an association) or cocaine CPP expression (recall of the association). The functional outcome of increased neuronal *Grin2c* expression in mPFC is also unknown. *Grin2c* encodes the NR2C subunit of the NMDA receptor, which is crucial for associative learning and gates long term plasticity^82^. *Grin2c* KO mice that lack NR2C subunit expression show an abnormal shift in E/I balance toward more inhibitory drive in layer V mPFC neurons, as well as a reduction in spine density and cognitive deficits^83^. Even a single acute exposure to cocaine has been shown to shift NMDA receptor subunit composition toward more NR3A-containing receptor expression^84^ which decreases calcium permeability and alters glutamatergic synaptic plasticity. Thus it will be critical to determine the functional consequences of a shift toward more NR2C-containing NMDA receptors in mPFC neurons in terms of both neuronal excitability and synaptic plasticity.

### Concluding remarks

Our study suggests neuronal *Grin2c* expression is a strong candidate component of the addicted engram since its expression is increased during cocaine-paired context-induced memory recall, and preventing its expression in neurons decreases cocaine CPP. While *Grin2c* was knocked down in all mPFC neurons to decrease CPP, Fos+ neurons expressed the majority of induced *Grin2c*, which makes it more likely that the knockdown effect was due to decreased *Grin2c* specifically in Fos-expressing neurons. Future studies using Fos-TRAP2 mice are required to confirm that *Grin2c* effects on behavior are due specifically to Fos-expressing neurons within ensembles mediating the CPP memory and thus strengthen the case for the role of *Grin2c* in the addicted engram.

## Acknowledgements

Plasmids were constructed and viral vectors were produced by the National Institute on Drug Abuse Genetic Engineering and Viral Vector Core Facility (RRID:SCR_022969).

## Authors’ contributions

L.A.R. and B.T.H. designed experiments and wrote the manuscript. The following authors performed experiments and/or data analysis: L.A.R., K.E.S., C.T.R., F.J.R., C.N.M., F.M.H., S.S.L., S.O., M.F.B., M.V., and R.M.; M.V. prepared the figures. All authors reviewed and approved the final manuscript.

---

**Figure S1.**
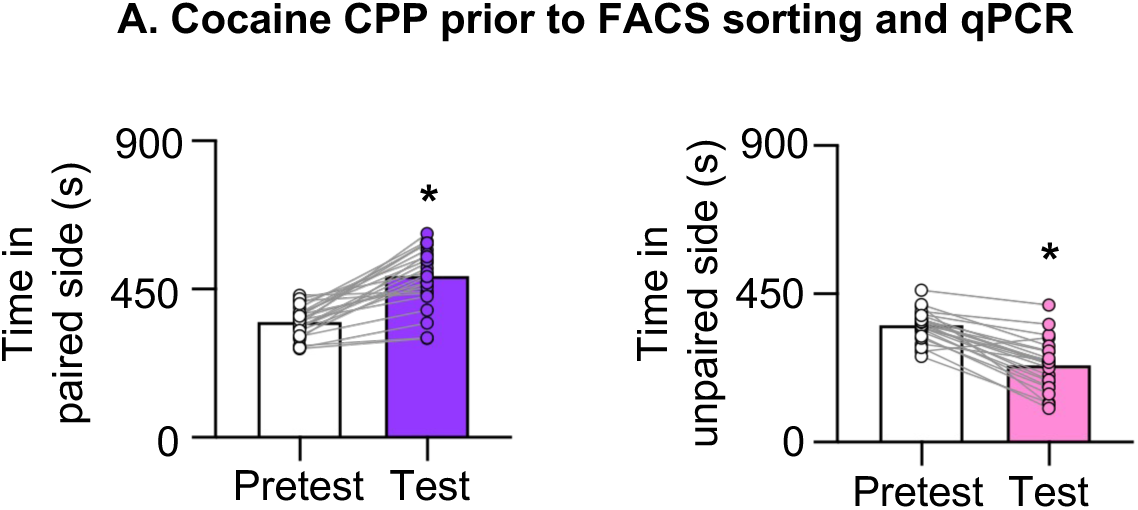
Cocaine conditioned place preference in mice used for FACS sorting and qPCR (related to. Figures 2 **and 3). (A)** Time (s) spent in the cocaine-paired (left) and unpaired (right) chambers at pretest versus test. Mice showed a significant increase in time spent in the paired chamber at test relative to pretest (T-test, [t_(23)_ = 8.66], *p < 0.05; n = 24), and a decrease in time spent in the unpaired chamber, (T-test, [t_(23)_ = 9.10], *p < 0.05; n = 24) confirming acquisition of cocaine CPP prior to FACS/qPCR analysis. Data are mean ± SEM; individual mice connected by lines.

**Figure S2.**
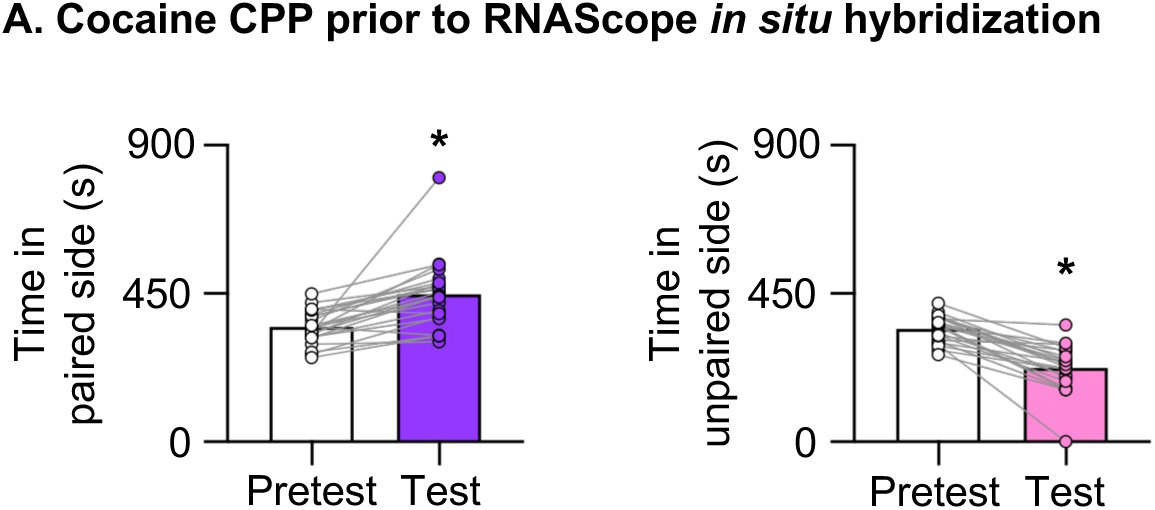
Cocaine conditioned place preference in mice used for RNAscope *in situ* hybridization (related to. Figure 4**). (A)** Time (s) spent in the cocaine-paired (left) and unpaired (right) chambers at pretest versus test. Mice showed a significant increase in time spent in the paired chamber at test relative to pretest (T-test, [t_(23)_ = 5.18], *p < 0.05; n = 24), and a decrease in time spent in the unpaired chamber, (T-test, [t_(23)_ = 9.11], *p < 0.05; n = 24) confirming acquisition of cocaine CPP prior to RNAscope analysis. Data are mean ± SEM; individual mice connected by lines.

