## Supplementary Table 1 for "Neuronal Grin2c expression and the cocaine engram: a medial prefrontal cortex substrate for conditioned place preference"

**Supplementary Table 1: Statistical analyses**

| **Figure number** | **Factor name** | **F and t values** | ***p*-value** |
| --- | --- | --- | --- |
| Figure 1B. (left) Cocaine CPP (Paired, seconds) | Test (within) | F _(1.6,24)_ = 11.78 | 0.0006 |
| Figure 1B. (right) Cocaine CPP (Unpaired, seconds) | Test (within) | F _(1.7,25)_ = 23.97 | <0.0001 |
| Figure 1C. (left) Cocaine CPP (Paired, percentage) | Test (within) | F _(1.6,24)_ = 11.78 | 0.0006 |
| Figure 1C. (right) Cocaine CPP (Unpaired, percentage) | Test (within) | F _(1.7,25)_ = 23.97 | <0.0001 |
| Figure 1D. Fos expression in PFC | Context (between) | F _(2,20)_ = 13.56 | 0.0002 |
| Figure 2D. (top left) NMDA receptor genes | Grin2c  Context (between)  Activity (within)  Context x Activity (mixed)  Grin2d  Context (between)  Activity (within)  Context x Activity (mixed)  Grin2b  Context (between)  Activity (within)  Context x Activity (mixed)  Grin1  Context (between)  Activity (within)  Context x Activity (mixed)  Grin2a  Context (between)  Activity (within)  Context x Activity (mixed) | F _(1,21)_ = 0.02  F _(1,21)_ = 12.52  F _(1,21)_ = 0.33  F _(1,28)_ = 3.12  F _(1,28)_ = 12.62  F _(1,28)_ = 0.38  F _(1,16)_ = 0.55  F _(1,13)_ = 6.29  F _(1,13)_ = 0.00  F _(1,16)_ = 2.47  F _(1,13)_ = 2.45  F _(1,13)_ = 0.20  F _(1,16)_ = 1.08  F _(1,13)_ = 1.29  F _(1,13)_ = 0.00 | 0.88  0.00  0.57  0.09  0.00  0.54  0.47  0.03  0.99  0.14  0.14  0.66  0.31  0.28  0.95 |
| Figure 2D. (top middle) Calcium related genes | Bdnf  Context (between)  Activity (within)  Context x Activity (mixed)  Fos  Context (between)  Activity (within)  Context x Activity (mixed)  ReIn  Context (between)  Activity (within)  Context x Activity (mixed)  CaMKII  Context (between)  Activity (within)  Context x Activity (mixed)  Creb1  Context (between)  Activity (within)  Context x Activity (mixed)  Cacna1c  Context (between)  Activity (within)  Context x Activity (mixed) | F _(1,15)_ = 1.00  F _(1,3)_ = 42.65  F _(1,3)_ = 0.03  F _(1,16)_ = 0.73  F _(1,12)_ = 10.04  F _(1,12)_ = 0.71  F _(1,16)_ = 0.05  F _(1,12)_ = 9.32  F _(1,12)_ = 0.30  F _(1,16)_ = 0.09  F _(1,12)_ = 26.68  F _(1,12)_ = 0.27  F _(1,28)_ = 3.81  F _(1,28)_ = 8.81  F _(1,28)_ = 2.01  F _(1,16)_ = 4.38  F _(1,13)_ = 15.06  F _(1,13)_ = 2.50 | 0.33  0.01  0.88  0.41  0.01  0.42  0.83  0.01  0.60  0.77  0.00  0.61  0.06  0.01  0.17  0.05  0.00  0.14 |
| Figure 2D. (top right) Scaffolding genes | Homer2  Context (between)  Activity (within)  Context x Activity (mixed)  Shank3  Context (between)  Activity (within)  Context x Activity (mixed)  Homer1  Context (between)  Activity (within)  Context x Activity (mixed)  Shank2  Context (between)  Activity (within)  Context x Activity (mixed)  Grip1  Context (between)  Activity (within)  Context x Activity (mixed)  Grip2  Context (between)  Activity (within)  Context x Activity (mixed) | F _(1,16)_ = 0.07  F _(1,9)_ = 11.14  F _(1,9)_ = 0.12  F _(1,28)_ = 1.52  F _(1,28)_ = 22.56  F _(1,28)_ = 0.06  F _(1,29)_ = 3.48  F _(1,29)_ = 14.09  F _(1,29)_ = 1.06  F _(1,16)_ = 0.32  F _(1,13)_ = 9.42  F _(1,13)_ = 0.02  F _(1,29)_ = 2.36  F _(1,29)_ = 4.91  F _(1,29)_ = 0.25  F _(1,27)_ = 2.47  F _(1,27)_ = 3.01  F _(1,27)_ = 0.04 | 0.80  0.01  0.74  0.23  <0.0001  0.81  0.07  0.001  0.31  0.58  0.01  0.89  0.14  0.03  0.62  0.13  0.09  0.84 |
| Figure 2D. (bottom left) AMPA/GABA receptor genes | Gabra1  Context (between)  Activity (within)  Context x Activity (mixed)  Gabbr1  Context (between)  Activity (within)  Context x Activity (mixed)  Gria3  Context (between)  Activity (within)  Context x Activity (mixed)  Gria1  Context (between)  Activity (within)  Context x Activity (mixed)  Gria2  Context (between)  Activity (within)  Context x Activity (mixed) | F _(1,16)_ = 1.21  F _(1,13)_ = 13.80  F _(1,13)_ = 0.47  F _(1,16)_ = 2.16  F _(1,13)_ = 10.93  F _(1,13)_ = 0.03  F _(1,16)_ = 1.95  F _(1,13)_ = 0.66  F _(1,13)_ = 1.72  F _(1,16)_ = 3.92  F _(1,13)_ = 4.56  F _(1,13)_ = 0.03  F _(1,16)_ = 2.84  F _(1,13)_ = 0.16  F _(1,13)_ = 0.01 | 0.29  0.00  0.51  0.16  0.01  0.86  0.18  0.43  0.21  0.07  0.05  0.86  0.11  0.70  0.91 |
| Figure 2D. (bottom middle) Potassium channel genes | Kcnj2  Context (between)  Activity (within)  Context x Activity (mixed)  Kcnn2  Context (between)  Activity (within)  Context x Activity (mixed)  Kcnip3  Context (between)  Activity (within)  Context x Activity (mixed)  Kcnma1  Context (between)  Activity (within)  Context x Activity (mixed)  Kcnn3  Context (between)  Activity (within)  Context x Activity (mixed) | F _(1,16)_ = 1.08  F _(1,6)_ = 4.69  F _(1,6)_ = 0.40  F _(1,15)_ = 0.05  F _(1,13)_ = 10.69  F _(1,13)_ = 0.57  F _(1,16)_ = 0.44  F _(1,13)_ = 7.34  F _(1,13)_ = 2.30  F _(1,29)_ = 0.50  F _(1,29)_ = 0.93  F _(1,29)_ = 0.00  F _(1,29)_ = 3.22  F _(1,29)_ = 1.88  F _(1,29)_ = 1.40 | 0.31  0.07  0.55  0.82  0.01  0.46  0.52  0.02  0.15  0.48  0.34  0.97  0.08  0.18  0.25 |
| Figure 2D. (bottom right) GPCR genes | Drd2  Context (between)  Activity (within)  Context x Activity (mixed)  Grm1  Context (between)  Activity (within)  Context x Activity (mixed)  Drd1  Context (between)  Activity (within)  Context x Activity (mixed)  Grm5  Context (between)  Activity (within)  Context x Activity (mixed) | F _(1,23)_ = 1.84  F _(1,23)_ = 0.08  F _(1,23)_ = 1.60  F _(1,16)_ = 0.02  F _(1,13)_ = 3.29  F _(1,13)_ = 0.06  F _(1,25)_ = 0.97  F _(1,25)_ = 1.97  F _(1,25)_ = 0.09  F _(1,16)_ = 1.53  F _(1,13)_ = 1.52  F _(1,13)_ = 0.07 | 0.19  0.79  0.22  0.88  0.09  0.81  0.34  0.17  0.77  0.23  0.24  0.79 |
| Figure 2B. qPCR of FACS sorted PFC tissue (all data heat map) | Gene (between)  Context (between)  Activity (within)  Gene x Context (between)  Gene x Activity (mixed)  Context x Activity (mixed)  Gene x Context x Activity (mixed) | F _(29,448)_ = 14.86  F _(1,448)_ = 1.93  F _(1,371)_ = 86.12  F _(29,448)_ = 1.00  F _(29,371)_ = 6.01  F _(1,371)_ = 0.27  F _(29,371)_ = 0.77 | <0.0001  0.17  <0.0001  0.47  <0.0001  0.60  0.80 |
| Figure 3A. qPCR results from all genes analyzed from FACS sorted PFC tissue | Grin2c  Context (between)  Activity (within)  Context x Activity (mixed)  Drd2  Context (between)  Activity (within)  Context x Activity (mixed)  Bdnf  Context (between)  Activity (within)  Context x Activity (mixed)  Fos  Context (between)  Activity (within)  Context x Activity (mixed)  Reln  Context (between)  Activity (within)  Context x Activity (mixed)  Gabra1  Context (between)  Activity (within)  Context x Activity (mixed)  CaMKII  Context (between)  Activity (within)  Context x Activity (mixed)  Kcnj2  Context (between)  Activity (within)  Context x Activity (mixed)  Homer2  Context (between)  Activity (within)  Context x Activity (mixed)  Shank3  Context (between)  Activity (within)  Context x Activity (mixed)  Grm1  Context (between)  Activity (within)  Context x Activity (mixed)  Homer1  Context (between)  Activity (within)  Context x Activity (mixed)  Kcnn2  Context (between)  Activity (within)  Context x Activity (mixed)  Shank2  Context (between)  Activity (within)  Context x Activity (mixed)  Gabbr1  Context (between)  Activity (within)  Context x Activity (mixed)  Grin2d  Context (between)  Activity (within)  Context x Activity (mixed)  Drd1  Context (between)  Activity (within)  Context x Activity (mixed)  Kcnip3  Context (between)  Activity (within)  Context x Activity (mixed)  Creb1  Context (between)  Activity (within)  Context x Activity (mixed)  Grin2b  Context (between)  Activity (within)  Context x Activity (mixed)  Grip1  Context (between)  Activity (within)  Context x Activity (mixed)  Cacna1c  Context (between)  Activity (within)  Context x Activity (mixed)  Gria3  Context (between)  Activity (within)  Context x Activity (mixed)  Kcnma1  Context (between)  Activity (within)  Context x Activity (mixed)  Grip2  Context (between)  Activity (within)  Context x Activity (mixed)  Gria1  Context (between)  Activity (within)  Context x Activity (mixed)  Grin1  Context (between)  Activity (within)  Context x Activity (mixed)  Grin2a  Context (between)  Activity (within)  Context x Activity (mixed)  Grm5  Context (between)  Activity (within)  Context x Activity (mixed)  Kcnn3  Context (between)  Activity (within)  Context x Activity (mixed)  Gria2  Context (between)  Activity (within)  Context x Activity (mixed) | F _(1,21)_ = 0.02  F _(1,21)_ = 12.52  F _(1,21)_ = 0.33  F _(1,23)_ = 1.84  F _(1,23)_ = 0.08  F _(1,23)_ = 1.60  F _(1,15)_ = 1.00  F _(1,3)_ = 42.65  F _(1,3)_ = 0.03  F _(1,16)_ = 0.73  F _(1,12)_ = 10.04  F _(1,12)_ = 0.71  F _(1,16)_ = 0.05  F _(1,12)_ = 9.32  F _(1,12)_ = 0.30  F _(1,16)_ = 1.21  F _(1,13)_ = 13.80  F _(1,13)_ = 0.47  F _(1,16)_ = 0.09  F _(1,12)_ = 26.68  F _(1,12)_ = 0.27  F _(1,16)_ = 1.08  F _(1,6)_ = 4.69  F _(1,6)_ = 0.40  F _(1,16)_ = 0.07  F _(1,9)_ = 11.14  F _(1,9)_ = 0.12  F _(1,28)_ = 1.52  F _(1,28)_ = 22.56  F _(1,28)_ = 0.06  F _(1,16)_ = 0.02  F _(1,13)_ = 3.29  F _(1,13)_ = 0.06  F _(1,29)_ = 3.48  F _(1,29)_ = 14.09  F _(1,29)_ = 1.06  F _(1,15)_ = 0.05  F _(1,13)_ = 10.69  F _(1,13)_ = 0.57  F _(1,16)_ = 0.32  F _(1,13)_ = 9.42  F _(1,13)_ = 0.02  F _(1,16)_ = 2.16  F _(1,13)_ = 10.93  F _(1,13)_ = 0.03  F _(1,28)_ = 3.12  F _(1,28)_ = 12.62  F _(1,28)_ = 0.38  F _(1,25)_ = 0.97  F _(1,25)_ = 1.97  F _(1,25)_ = 0.09  F _(1,16)_ = 0.44  F _(1,13)_ = 7.34  F _(1,13)_ = 2.30  F _(1,28)_ = 3.81  F _(1,28)_ = 8.81  F _(1,28)_ = 2.01  F _(1,16)_ = 0.55  F _(1,13)_ = 6.29  F _(1,13)_ = 0.00  F _(1,29)_ = 2.36  F _(1,29)_ = 4.91  F _(1,29)_ = 0.25  F _(1,16)_ = 4.38  F _(1,13)_ = 15.06  F _(1,13)_ = 2.50  F _(1,16)_ = 1.95  F _(1,13)_ = 0.66  F _(1,13)_ = 1.72  F _(1,29)_ = 0.50  F _(1,29)_ = 0.93  F _(1,29)_ = 0.00  F _(1,27)_ = 2.47  F _(1,27)_ = 3.01  F _(1,27)_ = 0.04  F _(1,16)_ = 3.92  F _(1,13)_ = 4.56  F _(1,13)_ = 0.03  F _(1,16)_ = 2.47  F _(1,13)_ = 2.45  F _(1,13)_ = 0.20  F _(1,16)_ = 1.08  F _(1,13)_ = 1.29  F _(1,13)_ = 0.00  F _(1,16)_ = 1.53  F _(1,13)_ = 1.52  F _(1,13)_ = 0.07  F _(1,29)_ = 3.22  F _(1,29)_ = 1.88  F _(1,29)_ = 1.40  F _(1,16)_ = 2.84  F _(1,13)_ = 0.16  F _(1,13)_ = 0.01 | 0.88  0.00  0.57  0.19  0.79  0.22  0.33  0.01  0.88  0.41  0.01  0.42  0.83  0.01  0.60  0.29  0.00  0.51  0.77  0.00  0.61  0.31  0.07  0.55  0.80  0.01  0.74  0.23  <0.0001  0.81  0.88  0.09  0.81  0.07  0.001  0.31  0.82  0.01  0.46  0.58  0.01  0.89  0.16  0.01  0.86  0.09  0.00  0.54  0.34  0.17  0.77  0.52  0.02  0.15  0.06  0.01  0.17  0.47  0.03  0.99  0.14  0.03  0.62  0.05  0.00  0.14  0.18  0.43  0.21  0.48  0.34  0.97  0.13  0.09  0.84  0.07  0.05  0.86  0.14  0.14  0.66  0.31  0.28  0.95  0.23  0.24  0.79  0.08  0.18  0.25  0.11  0.70  0.91 |
| Figure 3B. Activity-dependent genes | Grin2c  Context (between)  Activity (within)  Context x Activity (mixed)  Bdnf  Context (between)  Activity (within)  Context x Activity (mixed)  Fos  Context (between)  Activity (within)  Context x Activity (mixed)  ReIn  Context (between)  Activity (within)  Context x Activity (mixed)  Gabra1  Context (between)  Activity (within)  Context x Activity (mixed)  CaMKII  Context (between)  Activity (within)  Context x Activity (mixed)  Kcnj2  Context (between)  Activity (within)  Context x Activity (mixed)  Homer2  Context (between)  Activity (within)  Context x Activity (mixed)  Shank3  Context (between)  Activity (within)  Context x Activity (mixed)  Grm1  Context (between)  Activity (within)  Context x Activity (mixed)  Homer1  Context (between)  Activity (within)  Context x Activity (mixed)  Kcnn2  Context (between)  Activity (within)  Context x Activity (mixed)  Shank2  Context (between)  Activity (within)  Context x Activity (mixed)  Gabbr1  Context (between)  Activity (within)  Context x Activity (mixed)  Grin2d  Context (between)  Activity (within)  Context x Activity (mixed)  Kcnip3  Context (between)  Activity (within)  Context x Activity (mixed)  Creb1  Context (between)  Activity (within)  Context x Activity (mixed)  Grin2b  Context (between)  Activity (within)  Context x Activity (mixed)  Grip1  Context (between)  Activity (within)  Context x Activity (mixed)  Cacna1c  Context (between)  Activity (within)  Context x Activity (mixed)  Grip2  Context (between)  Activity (within)  Context x Activity (mixed)  Gria1  Context (between)  Activity (within)  Context x Activity (mixed) | F _(1,21)_ = 0.02  F _(1,21)_ = 12.52  F _(1,21)_ = 0.33  F _(1,15)_ = 1.00  F _(1,3)_ = 42.65  F _(1,3)_ = 0.03  F _(1,16)_ = 0.73  F _(1,12)_ = 10.04  F _(1,12)_ = 0.71  F _(1,16)_ = 0.05  F _(1,12)_ = 9.32  F _(1,12)_ = 0.30  F _(1,16)_ = 1.21  F _(1,13)_ = 13.80  F _(1,13)_ = 0.47  F _(1,16)_ = 0.09  F _(1,12)_ = 26.68  F _(1,12)_ = 0.27  F _(1,16)_ = 1.08  F _(1,6)_ = 4.69  F _(1,6)_ = 0.40  F _(1,16)_ = 0.07  F _(1,9)_ = 11.14  F _(1,9)_ = 0.12  F _(1,28)_ = 1.52  F _(1,28)_ = 22.56  F _(1,28)_ = 0.06  F _(1,16)_ = 0.02  F _(1,13)_ = 3.29  F _(1,13)_ = 0.06  F _(1,29)_ = 3.48  F _(1,29)_ = 14.09  F _(1,29)_ = 1.06  F _(1,15)_ = 0.05  F _(1,13)_ = 10.69  F _(1,13)_ = 0.57  F _(1,16)_ = 0.32  F _(1,13)_ = 9.42  F _(1,13)_ = 0.02  F _(1,16)_ = 2.16  F _(1,13)_ = 10.93  F _(1,13)_ = 0.03  F _(1,28)_ = 3.12  F _(1,28)_ = 12.62  F _(1,28)_ = 0.38  F _(1,16)_ = 0.44  F _(1,13)_ = 7.34  F _(1,13)_ = 2.30  F _(1,28)_ = 3.81  F _(1,28)_ = 8.81  F _(1,28)_ = 2.01  F _(1,16)_ = 0.55  F _(1,13)_ = 6.29  F _(1,13)_ = 0.00  F _(1,29)_ = 2.36  F _(1,29)_ = 4.91  F _(1,29)_ = 0.25  F _(1,16)_ = 4.38  F _(1,13)_ = 15.06  F _(1,13)_ = 2.50  F _(1,27)_ = 2.47  F _(1,27)_ = 3.01  F _(1,27)_ = 0.04  F _(1,16)_ = 3.92  F _(1,13)_ = 4.56  F _(1,13)_ = 0.03 | 0.88  0.00  0.57  0.33  0.01  0.88  0.41  0.01  0.42  0.83  0.01  0.60  0.29  0.00  0.51  0.77  0.00  0.61  0.31  0.07  0.55  0.80  0.01  0.74  0.23  <0.0001  0.81  0.88  0.09  0.81  0.07  0.001  0.31  0.82  0.01  0.46  0.58  0.01  0.89  0.16  0.01  0.86  0.09  0.00  0.54  0.52  0.02  0.15  0.06  0.01  0.17  0.47  0.03  0.99  0.14  0.03  0.62  0.05  0.00  0.14  0.13  0.09  0.84  0.07  0.05  0.86 |
| Figure 3C. Context-dependent genes | Homer1  Context (between)  Activity (within)  Context x Activity (mixed)  Grin2d  Context (between)  Activity (within)  Context x Activity (mixed)  Creb1  Context (between)  Activity (within)  Context x Activity (mixed)  Cacna1c  Context (between)  Activity (within)  Context x Activity (mixed)  Gria1  Context (between)  Activity (within)  Context x Activity (mixed)  Kcnn3  Context (between)  Activity (within)  Context x Activity (mixed) | F _(1,29)_ = 3.48  F _(1,29)_ = 14.09  F _(1,29)_ = 1.06  F _(1,28)_ = 3.12  F _(1,28)_ = 12.62  F _(1,28)_ = 0.38  F _(1,28)_ = 3.81  F _(1,28)_ = 8.81  F _(1,28)_ = 2.01  F _(1,16)_ = 4.38  F _(1,13)_ = 15.06  F _(1,13)_ = 2.50  F _(1,16)_ = 3.92  F _(1,13)_ = 4.56  F _(1,13)_ = 0.03  F _(1,29)_ = 3.22  F _(1,29)_ = 1.88  F _(1,29)_ = 1.40 | 0.07  0.001  0.31  0.09  0.00  0.54  0.06  0.01  0.17  0.05  0.00  0.14  0.07  0.05  0.86  0.08  0.18  0.25 |
| Figure 4C. (top left) NeuN-negative Grin2c cells | Context (between) | F _(2,13)_ = 1.07 | 0.37 |
| Figure 4C. (top right) NeuN-positive Grin2c cells | Context (between) | F _(2,13)_ = 23.00 | <0.0001 |
| Figure 4C. (bottom left) Fos-negative Grin2c cells | Context (between) | F _(2,13)_ = 1.09 | 0.37 |
| Figure 4C. (bottom right) Fos-positive Grin2c cells | Context (between) | F _(2,13)_ = 4.90 | 0.03 |
| Figure 4D. Grin2c cells in PFC: total | Context (between) | F _(2,13)_ = 0.42 | 0.66 |
| Figure 5D.  Virus Expression | Virus (between)  NeuN (within)  NeuN x Virus | F _(1,13)_ = 0.72  F _(1,13)_ = 38.44  F _(1,13)_ = 1.19 | 0.41  <0.0001  0.30 |
| Figure 5F. Cocaine CPP after viral injection in PFC | Virus (between)  Test (within)  Test x Virus | F _(2,18)_ = 4.30  F _(1.40,25.20)_ = 6.72  F _(2.80,25.20)_ = 0.21 | 0.03  0.01  0.88 |
| Figure S1. (left)  Cocaine CPP prior to FACS sorting and qPCR (Paired, seconds) |  | T _(23)_ = 8.66 | <0.0001 |
| Figure S1. (right)  Cocaine CPP prior to FACS sorting and qPCR (Unpaired, seconds) |  | T _(23)_ = 9.10 | <0.0001 |
| Figure S2. (left)  Cocaine CPP prior to RNAScope in situ hybridization (Paired, seconds) |  | T _(23)_ = 5.18 | <0.0001 |
| Figure S2. (right)  Cocaine CPP prior to RNAScope in situ hybridization (Unpaired, seconds) |  | T _(23)_ = 9.11 | <0.0001 |
