## Supplementary Table 2 for "Neuronal Grin2c expression and the cocaine engram: a medial prefrontal cortex substrate for conditioned place preference"

**Supplementary Table 2. Primer/probe table for genes analyzed by qPCR**

| Gene | TaqMan probe | Forward primer | Reverse primer |
| --- | --- | --- | --- |
| *Arc* | Mm01204954_g1* |  |  |
| *Bdnf* | AAGCCACAATGTTCCACCAG | CACATTACCTTCCTGCATCTGTTG | ACCATAGTAAGGAAAAGGATGGTCAT |
| *Cacna1c* | Mm01188822_m1* |  |  |
| *Creb1* | Mm00501607_m1* |  |  |
| *Drd1* | Mm01353211_m1* |  |  |
| *Drd2* | Mm00438545_m1* |  |  |
| *Fos* | Mm00487425_m1* |  |  |
| *Gabbr1* | Mm00444578_m1* |  |  |
| *Gabra1* | Mm00439046_m1* |  |  |
| *Gria1* | Mm00433753_m1* |  |  |
| *Gria2* | Mm00442822_m1* |  |  |
| *Gria3* | Mm00497506_m1* |  |  |
| *Grin1* | Mm00433790_m1* |  |  |
| *Grin2a* | Mm00433802_m1* |  |  |
| *Grin2b* | Mm00433820_m1* |  |  |
| *Grin2c* | Mm00439180_m1* |  |  |
| *Grin2d* | Mm00433822_m1* |  |  |
| *Grip1* | Mm00503847_m1* |  |  |
| *Grip2* | Mm01183472_m1* |  |  |
| *Grm1* | Mm00810219_m1* |  |  |
| *Grm5* | Mm00690332_m1* |  |  |
| *Homer1* | Mm00516275_m1* |  |  |
| *Homer2* | Mm01314936_m1* |  |  |
| *Kcnip3* | Mm01339777_m1* |  |  |
| *Kcnj2* | Mm00434616_m1* |  |  |
| *Kcnma1* | Mm01268569_m1* |  |  |
| *Kcnn2* | Mm00446514_m1* |  |  |
| *Kcnn3* | Mm00446516_m1* |  |  |
| *RbFox3* | Mm01248771_m1* |  |  |
| *Reln* | Mm00465200_m1* |  |  |
| *Shank2* | Mm00683065_m1* |  |  |
| *Shank3* | Mm00498775_m1* |  |  |

* TaqMan catalog number for ordering from Applied Biosystems.
